# Enhanced multisensory integration in the olfactory bulb of the Mexican cavefish

**DOI:** 10.64898/2026.02.26.708145

**Authors:** Evan Lloyd, Anna Koga, Douglas A. Storace

**Affiliations:** Department of Biological Science, Florida State University, Tallahassee, FL; Program in Neuroscience, Florida State University, Tallahassee, FL; Institute of Molecular Biophysics, Florida State University, Tallahassee, FL

## Abstract

*Astyanax mexicanus* consists of eyed, river-dwelling “surface” fish, and multiple, independently evolved cave populations, which have converged on troglobitic traits such as eye loss and reduced metabolism. However, considerably less is known about constructive adaptations, which include a larger olfactory epithelium in cavefish. It is unknown how this relates to the olfactory bulb (OB), which is the first stage of olfactory sensory processing in the brain. The goal of the present study is to begin to define the structure and functional organization of the OB in *A. mexicanus*, and to begin to understand how it was transformed via cave adaptation. We addressed these questions using whole-mount immunohistochemistry and *in vivo* Ca^2+^ imaging from the OB of developmentally matched surface and Pachón cavefish. The cavefish OB was significantly larger than surface fish by 14 days post fertilization (dpf), which was accompanied by a broad and proportional increase in synaptic input to most glomerular regions. Increases in the size of the OB were accompanied by increases in the number of neurons expressing tyrosine hydroxylase and calretinin, the latter of which occurred primarily in the medial OB and could not be explained as a compensatory response to a larger OB. *In vivo* Ca^2+^ imaging from the dorsal OB of surface and cavefish in response to a panel of chemical stimuli revealed odor-evoked responses that were spatially organized and highly conserved across the two populations. Surprisingly, the medial OB was consistently activated by any change in water flow in both populations, although the number of water-responsive neurons was significantly greater in cavefish when measurements were performed using either *in vivo* imaging or the neuronal activity marker phospho-ERK. Water-responding neurons were similarly present in the olfactory epithelium in both populations, along with neurons expressing the mechanosensitive ion channel Piezo2, with significantly more Piezo2-expressing neurons present in cavefish. Therefore, cavefish exhibit enhanced multisensory integration of olfactory and mechanosensory input in the earliest stage of olfactory sensory processing in the brain.

## Introduction

The Mexican tetra *Astyanax mexicanus* exists as both a river-dwelling “surface” form, and multiple populations of cave-adapted fish which have converged on troglobitic phenotypes, including a prominent loss of vision and changes in metabolism. Consequently, comparative studies using the two forms have become an increasingly powerful approach to understand the role of evolution-mediated changes in brain structure and function. However, surprisingly little is understood about potentially constructive changes that can occur as the result of adaptation to a cave environment. One example of this includes a larger peripheral olfactory organ present in cavefish (Hinaux et al., 2016; Kuball et al., 2024). Because fish use their olfactory system to detect food and mates and avoid predators, changes to their olfactory processing abilities may reflect important behaviorally relevant adaptations to their unique trophic environment (Edmunds et al., 2016; Keagy et al., 2018; Crowley-Gall et al., 2019; Álvarez-Ocaña et al., 2023).

In vertebrate fish species, olfactory detection begins with chemosensory stimuli entering the naris which activate olfactory sensory neurons (OSNs) in the olfactory epithelium (OE). The OSNs project into the olfactory bulb (OB), where they synapse into glomeruli, regions of neuropil which receive convergent input from specific odor receptors in most vertebrate species (Kermen et al., 2013; Olivares and Schmachtenberg, 2019). Different classes of odor broadly activate different regions across the OB, creating a chemotopic mapping from the epithelium onto the OB (Friedrich and Korsching, 1997, 1998). Glomeruli are surrounded by populations of interneurons and projection neurons, which are involved in the processing and transmission of the incoming OSN input to the rest of the brain (Miyasaka et al., 2013; Zhu et al., 2013; Wanner et al., 2016). Therefore, the OB is an obligatory interface between the peripheral olfactory organ and the rest of the brain. Surprisingly, no data exists characterizing the OB in *A. mexicanus* in general, and whether there are structural or functional differences between the two populations. We addressed these knowledge gaps using a combination of whole-mount immunohistochemistry at different days post fertilization (dpf), and *in vivo* Ca^2+^ imaging from the OB of transgenic surface and Pachón cavefish that express GCaMP6s pan-neuronally (Stahl et al., 2019; Jaggard et al., 2020; Lloyd et al., 2022).

Both surface and cavefish exhibited a rapid developmental increase in the size of the OB beginning at 21dpf. The projections from the OE to the OB innervated glomerular regions of interest that appeared conserved with other vertebrate fish species (Braubach et al., 2012; Braubach et al., 2013; Braubach and Croll, 2021). Notably, the cavefish OB was significantly larger than in the surface morphs beginning at 14dpf, and this was accompanied by broad increases in glomerular volume that were proportional to the size of the OB. We confirmed that neurons expressing the enzyme tyrosine hydroxylase (TH) and the protein calretinin (CR) were present in the OB of both morphs. Both surface and cavefish exhibited a significant increase in the number of TH-expressing neurons across development, with greater numbers present in cavefish. The numbers of CR-expressing neurons increased in cavefish across development but did not change in surface fish.

We performed *in vivo* Ca^2+^ imaging from the dorsal surface of the OB in transgenic surface and cavefish that express GCaMP6s pan-neuronally. A panel of liquid odor stimuli confirmed the presence of OB neurons activated by specific odors, which exhibited a spatial organization that was conserved within and across both populations. Surprisingly, our control water stimulus consistently activated the medial OB in both surface and cavefish, indicating that OB neurons receive sensory signals about changes in water flow. The number of water-responsive OB neurons was significantly greater in cavefish when measured using *in vivo* Ca^2+^ imaging and when measuring the number of OB neurons expressing the neuronal activity marker phosphorylated ERK after water stimulation. The olfactory epithelium in both populations contained neurons that were water-responsive, and which expressed the mechanosensitive ion channel Piezo2. Remarkably, the number of Piezo2-expressing neurons in the OE was significantly greater in the cave population.

Our results provide the first characterization of the structural and functional organization of the OB in *A. mexicanus*, revealing both broadly conserved properties and unique adaptations. Surprisingly, the larger cavefish OB appears to reflect proportional increases in sensory input to different glomeruli, as opposed to selective changes in specific glomeruli. The presence of water-responsive neurons and Piezo2-expressing neurons in the olfactory epithelium suggests that the OB of both surface and cavefish inherits and integrates both olfactory and mechanosensitive stimuli from the periphery. Higher levels of Piezo2-expressing neurons present in the cavefish epithelium indicate that cavefish have potentially enhanced multisensory integration beginning at the first stage of olfactory sensory processing in the brain.

## Methods

| Reagent | Source | Item No. |
| --- | --- | --- |
| **Histology** |  |  |
| Glyoxal (40% solution) | Sigma-Aldrich | 128465 |
| Acetic acid | Sigma-Aldrich | A6283 |
| Sodium Hydroxide (NaOH) | Sigma-Aldrich | 221456 |
| Phosphate Buffered Saline (PBS), 10x | ThermoFisher | BP399-1 |
| Triton X-100 | MP Biomedicals | 04807426-CF |
| Dimethyl Sulfoxide (DMSO) | Ward's Science | 470300-982 |
| Low Melting Point (LMP) Agarose | IBI Scientific | IB70056 |
| **Antibodies** |  |  |
| Mouse anti-SV2 | Developmental Studies Hybridoma Bank | SV2 |
| Rabbit anti-TH | Sigma-Aldrich | AB152 |
| Rabbit anti-calretinin | Swant | CR7697 |
| Mouse anti-Phospho-p44/42 MAPK (pERK) | Cell Signaling Technology | 5726VW |
| Mouse anti-Piezo2 | ThermoFisher | PA5-72975 |
| NeuroTrace 435/455 | ThermoFisher | N21479 |
| iFluor 555 goat anti-mouse IgG (H+L) | AAT Bioquest | 16776 |
| iFluor 647 goat anti-rabbit IgG (H+L) | AAT Bioquest | 16837 |
| **Odorants** |  |  |
| Sodium Chloride (NaCl) | Avantor | 97063-366 |
| Lithocholic Acid (LCA) | Sigma-Aldrich | L6250 |
| Adenosine | Sigma-Aldrich | 01890 |
| L-Alanine | Sigma-Aldrich | A7469 |
| Cadaverine-dihydrochloride | Sigma-Aldrich | 33220 |
| Ammonium Chloride (NH<sub>4</sub>Cl) | Sigma-Aldrich | 11209 |
| **Imaging** |  |  |
| Mivacurium Chloride | AstaTech | G12763 |

### Animal husbandry

All procedures were approved by the Florida State University Animal Care and Use Committee. Fish used in all experiments were generated and raised at Florida State University in a dedicated aquarium facility. The aquarium temperature was maintained at 23°C on a 14 hour light:10 hour dark schedule as previously described (Kozol et al., 2023). Transgenic fish used for Ca^2+^ imaging were Tg(elav3:H2B-GCaMP6s) surface and cavefish, derived from previously generated lines (Jaggard et al., 2020; Lloyd et al., 2022). Non-transgenic surface fish used for immunohistochemistry (7-28dpf) were derived from fish collected at the Nacimiento del Rio Choy (San Luis Potosi, Mexico). Cavefish descended from inhabitants of the Pachón cave (Sierra del Abra, Mexico).

To stimulate breeding, water temperature was raised by ∼2°C with a submersible heater, and fish were fed flake food (TetraMin Pro) to satiation 2–3 times daily (Elipot et al., 2014). Following breeding, embryos were collected from tanks and raised in 2L aquaria at a density of ∼24 fish/L under the same temperature and light cycle. Starting at 7dpf, fish were fed live *Artemia* nauplii to satiation once daily, with >75% water changes 1x/week, until used in experiments. For both immunohistochemical and live imaging experiments, all fish were tested prior to daily feeding.

### Immunohistochemistry and confocal imaging

Surface and cavefish at each studied time point were euthanized by immersion in ice-cold (4°C) aquarium water, then immersed in ice cold glyoxal solution (9% glyoxal, 8% acetic acid, pH-adjusted to 4-5 with 5N NaOH solution) and fixed overnight at 4°C (Konno et al., 2023). Samples of both surface and cavefish at 28dpf were dissected out following fixation due to higher levels of pigmentation present in surface fish at that age. Fixed whole-mount samples were rinsed 3 times for 10 minutes each in ice-cold PBS with 0.1% Triton-X (PBS-T), followed by incubation in primary solution (PBS-T, with 1% DMSO, primary antibodies at 1:500 dilution) overnight at 4°C.

A subset of samples were labeled using NeuroTrace (**Figure 7F**), which was added along with the primary antibodies at a 1:100 dilution. Samples were rinsed for 3×10 minutes with ice-cold PBS-T, and then were incubated in secondary solution (PBS-T, with 1% DMSO, secondary antibodies at 1:500 dilution) overnight at 4°C. Following a final 3×10 minutes set of rinses in PBS-T, samples were mounted in 2% low-melting point (LMP) agarose for imaging. Whole-mount samples were imaged on a Nikon CSU-W1 spinning disk confocal using a 20x (NA 0.95) or 40x (NA 1.15) objective lens. The entire OB of each sample was imaged using Z-steps of 2 µm. Image brightness and contrast were uniformly adjusted in ImageJ and cropped and arranged in Adobe Illustrator.

### *In vivo* imaging and olfactometry

Surface and cavefish between 9-22dpf were paralyzed by immersion in 0.05% mivacurium chloride (AstaTech) and were mounted in LMP agarose in a Siskiyou perfusion chamber (Automate Scientific, PC-H). The agarose around the olfactory epithelia was carefully cleared and the sample was placed underneath an Olympus BX50WI microscope equipped with an LED (Prizmatix), optimal filter sets, a Nikon 16× 0.8 N.A. objective lens, and a CCD camera (NeuroCCD-SM256 RedShirtImaging, Decatur GA).

The microscope was equipped with an odor delivery system consisting of 8 independent syringe reservoirs filled with aquarium water, or liquid odors dissolved in aquarium water. Teflon tubing (NResearch, TBGM108) connected each reservoir to an independent solenoid valve (Part #1303, 3 msec switching duration) which was controlled using an automated valve controller (Sanworks) and custom-written software (MATLAB, Mathworks). The valve connected to an 8-channel perfusion pencil (Automate Scientific, 04-08-zdv) which was fit with a 250 µm tip (Automate Scientific, 04-250). The perfusion pencil was mounted to a 3-way micro-manipulator and was positioned in front of the nares of the sample. Aquarium water was continuously perfused over the subject at a rate of 300 µl min^-1^ throughout the entire experiment and was collected from the opposite end of the chamber via vacuum suction.

Imaging data were recorded at a spatial and temporal resolution of 256×256 pixels at 25 Hz. Olfactometer stimulus timing was synchronized to image acquisition with a National Instruments multifunction I/O device. Except for the trials involving repeated switching of water lines (**Figure 6**), individual imaging trials were 24 seconds in duration and included a 3-second baseline period, followed by 5 seconds of odor stimulation and a 16 second post-stimulus period. Switching experiment trials were performed similarly except that the water line was alternated every 5 seconds (0.1 Hz). The response to each odor stimulus was measured in 3 separate trials in each preparation. The mean of the individual trials was used for all data analysis.

Aquarium water was composed of de-ionized water with aquarium salt (Instant Ocean) added to bring the conductivity to ∼500 µS/cm, and the pH adjusted to 7 with the addition of sodium bicarbonate. With the exception of LCA, all odorants were dissolved in aquarium water and stored in stock solutions in concentrations of ≤1M at -20°C until the day of the experiment, at which point they were defrosted and diluted further in aquarium water to their working concentrations. LCA was dissolved in 100% DMSO at a concentration of 1M, then diluted in aquarium water to a stock concentration of 10^-3^M and stored at 4°C until the day of the experiment. All odorants were presented at a final concentration of 10^-4^M, except NaCl, which was presented at 10^-3^M. Odorants were never re-frozen, and excess dilutions were disposed of at the end of the experiment day.

### Water stimulation and pERK expression experiments

Samples used for the pERK expression analysis were prepared for *in vivo* imaging. Each sample in the “stimulus” group was prepared simultaneously with a paired “control” subject which was treated identically, except that the control subject did not have the agarose cleared from its epithelia. Following mounting in LMP agarose, the chamber containing both subjects was placed under the microscope, the perfusion pencil was aligned to the experimental subject, and stimulation was performed as follows: for a total of 5 minutes, water was perfused over the subject from two separate lines, alternating lines every 5 seconds. 2-3 minutes of each experiment was recorded to confirm stimulation of the subject. Immediately following the completion of the stimulation protocol, both the experimental and control subject were quickly removed from agarose, euthanized by immersion in ice-cold water, then immediately fixed overnight in glyoxal solution at 4°C. Following fixation, samples were treated as described in the IHC methods.

### Data analysis

#### IHC analysis

OB volume measurements and cell counts were performed in FIJI/ImageJ 1.54g (NIH). OB glomeruli were identified based on the presence of dense spherical volumes of SV2 expression and were differentiated by small boundaries lacking SV2 expression. Total OB volume and glomerular volume measurements were performed using the “Segmentation Editor” plug-in, using the freehand selection tool and the interpolation tool to outline the regions of interest. The resulting segmentations were imported into the 3DSuite plug-in (Ollion et al., 2013) and the total volume of each region of interest was extracted. For total OB volume, the sum of both OBs was reported. For glomerular volume measurements and for olfactory nerve diameter, both the left and right OBs were measured, and the average of the two measurements was reported. Cell counts were performed using the “Multi-point” tool; the center of each neuron was marked, and the total number of cells within both olfactory bulbs was reported. The numbers of Piezo2-expressing cells in the OE were quantified in a single nare in each sample (**Figure 7**).

#### Frame Subtraction Analysis

The frame subtractions analyses in **Figure 4** were generated using Turbo-SM software (SciMeasure, Decatur, GA). Intensity ranges are scaled equally across all odor responses within each subject, to highlight the spatial characteristics of the response. The mean fluorescence and frame subtraction images were generated from the average of three individual trials. The frame subtraction images were generated by subtracting an average of 10 frames during the odor stimulation period from the average of 10 frames preceding the odor.

#### Single-cell analysis

Regions of interest (ROIs) of single cells were generated manually by selecting the center point of each cell within a single olfactory bulb that produced a detectable response to any stimulus using Turbo-SM software. ROIs were manually adjusted between trials to compensate for slight drift of the subject in the XY plane when necessary. Any subjects which exhibited significant drift in the Z dimension were discarded from analysis, and individual trials with gross movement artifacts were discarded from subsequent analysis. The fluorescence time course from the ROIs were converted to ΔF/F by dividing the entire trace by the average of the 10 frames prior to stimulus onset, and were low-pass filtered at 5 Hz (**Figure 4C,5A, 5B**). Odor responses were defined as the mean ΔF/F of the first 30 frames of odor stimulation subtracted from the frames immediately preceding the stimulus. A response was considered significant if the peak of the ΔF/F signal during the stimulus was at least 6 standard deviations above the pre-stimulus period.

Lifetime sparseness (LS) was calculated as follows, according to the formula reported in (Zak et al., 2024):

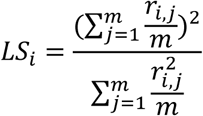

Where ***m*** = the number of stimuli and ***r_i,j_*** = the response of a cell ***i*** to a stimulus ***j***.

### Statistical Methods

All anatomical comparisons, including OB volume, ON nerve diameter, absolute glomerular volume, normalized glomerular volume, absolute cell counts, and normalized cell counts (**Figure 1-3**) were performed using multiple unpaired t-tests with Holm-Sidak correction for multiple comparisons. Comparisons of functional responses to olfactory stimuli were performed using 2-way repeated measures ANOVA with Sidak’s multiple comparisons *post-hoc* test to detect differences between populations. Comparison of Piezo2-expressing cells was performed using an unpaired t-test (**Figure 7E)**. All statistical comparisons were performed in Graphpad Prism (v. 10.6.0). All error bars throughout the manuscript indicate S.E.M.

**Figure 1:**
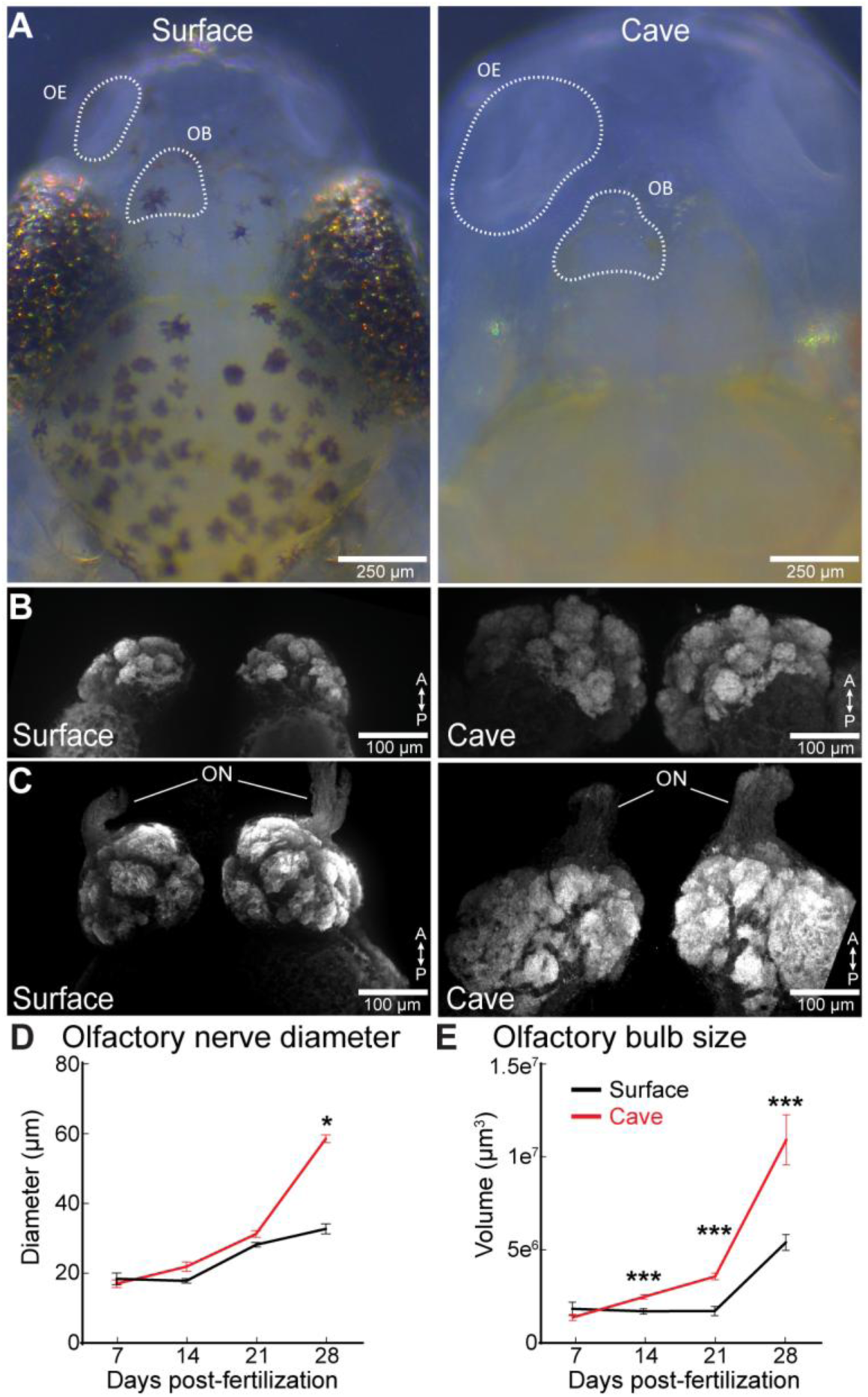
(A) Brightfield images of surface and cavefish at 28 days post-fertilization (dpf). The olfactory epithelium (OE) and olfactory bulb (OB) are outlined with dashed contours. (B-C) Z-projection of the surface and cavefish dorsal (B) and ventral (C) OB labeled with SV2. (D) Olfactory nerve (ON) diameter at 7, 14, 21 and 28dpf. (E) Volume of the surface and cave OB at 7,14,21,and 28dpf. * p<0.05; *** p<0.001.

## Results

### Structure of the olfactory bulb and olfactory nerve in A. mexicanus across development

Low magnification images from *A. mexicanus* indicate that both the olfactory epithelium (OE) and the olfactory bulb (OB) of the blind cavefish morphs are larger than those of surface morphs at 28 days post fertilization (dpf) (**Figure 1A***, OB*). We mapped the projections from the OE to the OB in cave and surface fish using immunohistochemistry for the synaptic vesicle marker SV2, which is enriched in axon projections (**Figure 1B-C**). Whole mount confocal imaging revealed that the olfactory nerve (ON) diameter and the OB size increased across development from 7 to 28dpf in both surface and cavefish (**Figure 1B-C**). However, the cavefish ON and OB were both significantly larger than in the surface fish by 28dpf (**Figure 1D, E**).

### The A. mexicanus OB contains stereotyped glomerular regions of interest that increase proportionally with OB volume

The ON formed 7 well defined glomerular regions of interest in the OB by 7dpf in both surface and cavefish whose structure and position were consistent with those in *D. rerio* (**Figure 2A**) (Braubach et al., 2012; Braubach et al., 2013). The dorsolateral (dlG), lateral (lG), medial anterior (maG), and ventromedial (vmG) glomerular regions were the largest regions, followed by the mediodorsal (mdG), the dorsal (dG) and the ventroposterior (vpG) (**Figure 2A**). Each glomerular region significantly increased in volume from 7 to 28dpf, and 6 of 7 were larger in cavefish than surface fish by 28 dpf (**Figure 2C**). However, normalizing the volume of each glomerulus to the total OB volume revealed that the size of each glomerulus was the same proportion of the OB at all developmental time points in both surface and cavefish (**Figure 2D**). Therefore, the larger OB volume in cavefish reflects a broad expansion of all glomeruli.

**Figure 2:**
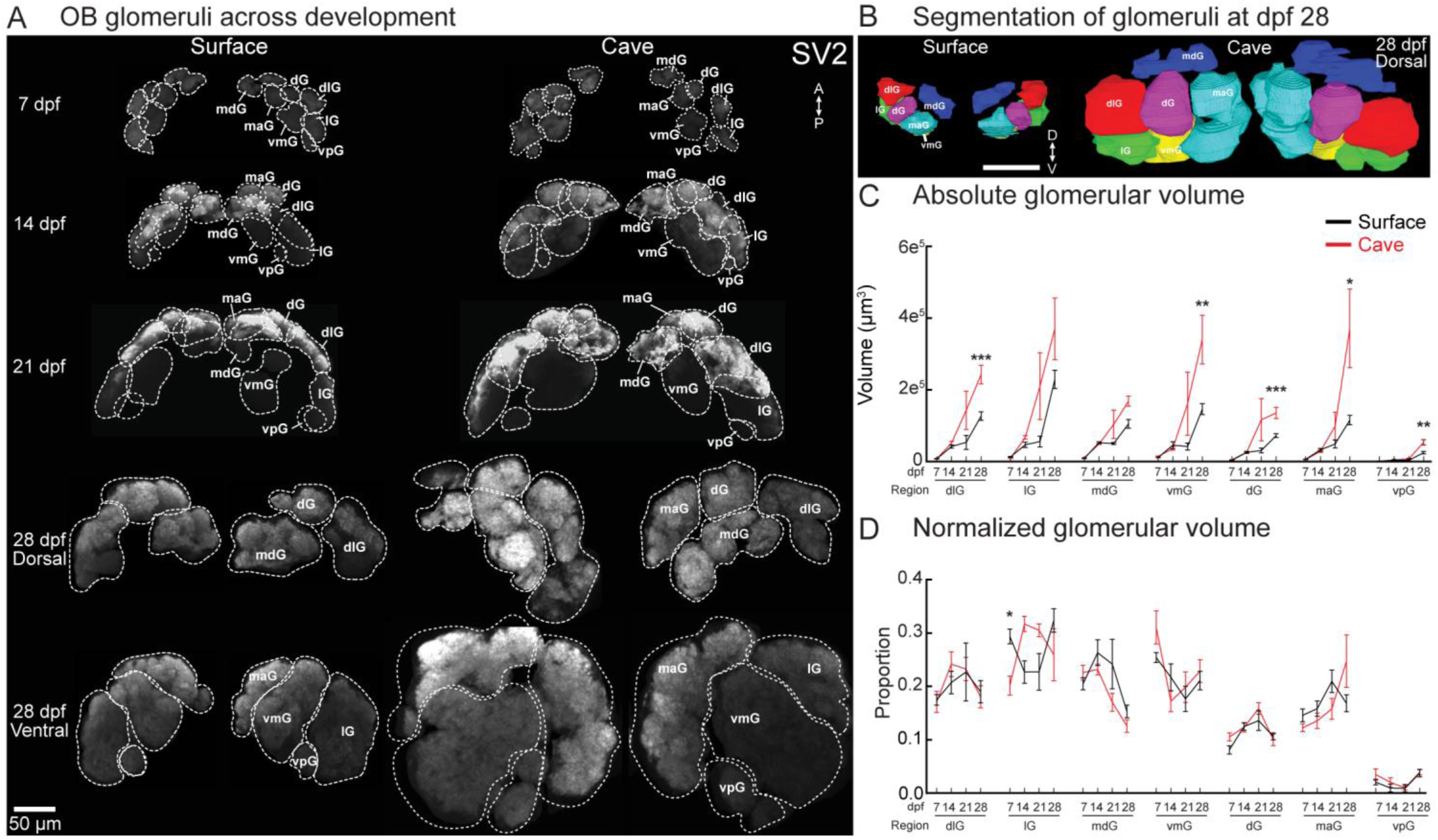
SV2 expression in the OB of surface and cavefish illustrating development of glomerular structure. (**A**) Z-projections of surface and cave OBs at 7, 14, 21, and 28dpf. The 28dpf time point is illustrated from both dorsal and ventral views. Contours indicate anatomically distinct glomeruli. All images are scaled to the same size. (**B**) Frontal (top) and dorsal (bottom) views of the segmentation of a surface and cavefish OB at 28dpf. (**C**) Volume of the glomerular regions at 7,14,21, and 28dpf in surface and cavefish. (**D**) The proportion of each glomerular region as a share of total glomerular volume at each time point. * indicates *p*<0.05, ** indicates *p*<0.01, *** indicates *p*<0.001. mdG, mediodorsal glomeruli; dG, dorsal glomeruli; dlG, dorsolateral glomeruli; lG, lateral glomeruli; maG, medioanterior glomeruli; vmG, ventromedial glomeruli; vpG, ventroposterior glomeruli.

### The numbers of TH-expressing neurons increase proportionally with OB volume, while calretinin-expressing neurons are selectively upregulated in cavefish

The vertebrate OB contains a complex synaptic network comprised of multiple cell types that include neurons that express the dopamine precursor tyrosine hydroxylase (TH) and the calcium binding protein calretinin (Castro et al., 2006; Parrish-Aungst et al., 2007; Bundschuh et al., 2012; Friedrich, 2013; Zhu et al., 2013). We performed whole-mount immunohistochemistry to confirm that both cell types were present in the OB of both surface and cavefish (**Figure 3A-B**). TH-expressing neurons were widespread in both the glomerular layer and the mitral cell layer of both surface and cavefish by 28dpf (**Figure 3A**). The numbers of TH-expressing neurons increased across development in both morphs, with cavefish having significantly more neurons at 14, 21 and 28dpf (**Figure 3C**). Normalizing the numbers of TH-expressing neurons to OB volume revealed that except for a small difference at 14dpf, these increases occurred proportionally as a function of overall synaptic volume (**Figure 3C**, right).

**Figure 3:**
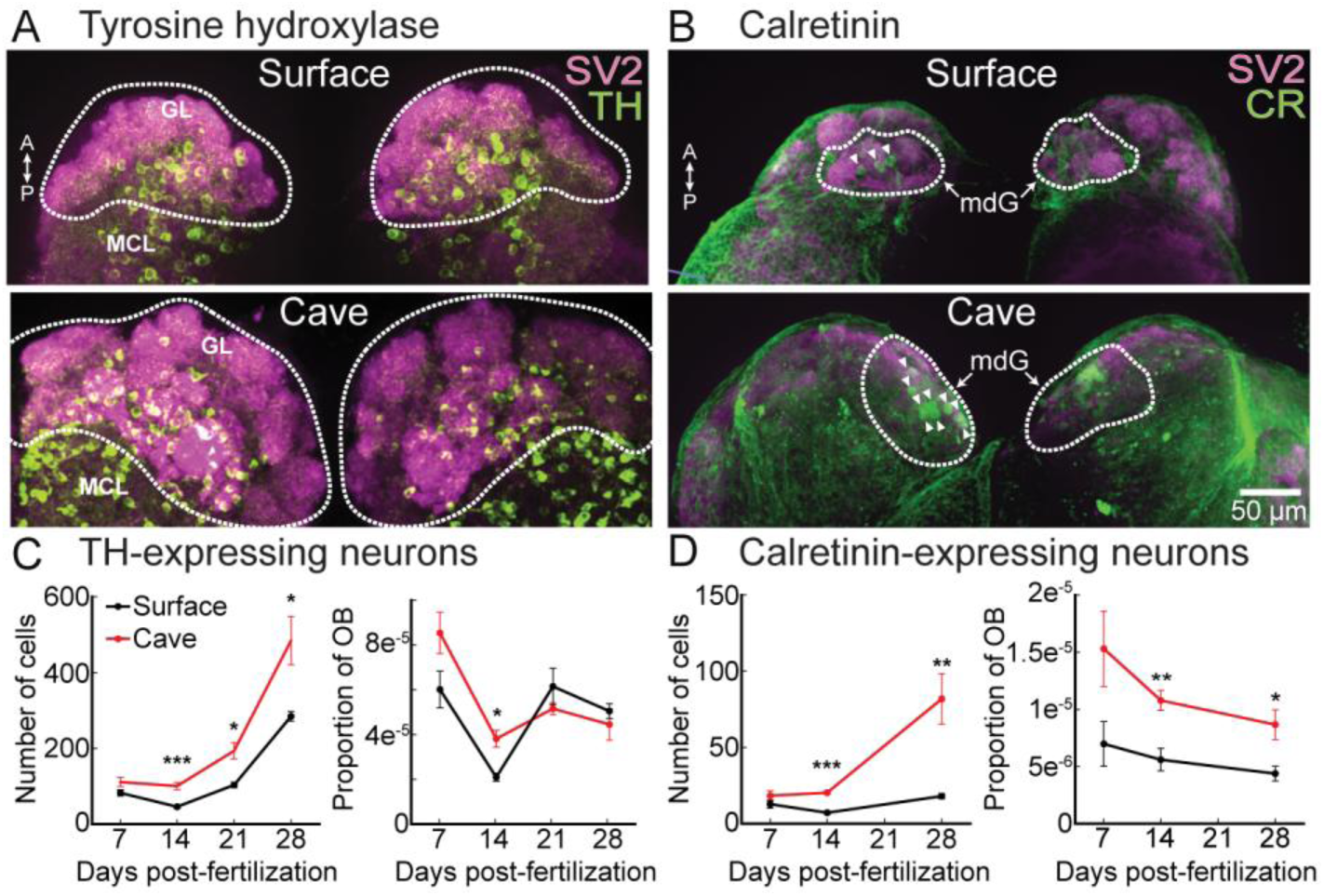
(A-B) Tyrosine hydroxylase (TH) (A) and calretinin (B) expression in the surface (top) and cavefish (bottom) OB. (C-D) TH-(C) and calretinin-expressing (D) neurons expressed as cell counts (left) and as a proportion of the OB volume (right). GL, glomerular layer; MCL, mitral cell layer, mdG, mediodorsal glomeruli. Arrowheads in (B) denote calretinin-positive neurons.

In contrast, calretinin neurons were more spatially localized around the mdG glomerular region, with sparser expression elsewhere throughout the OB (**Figure 3B**). Surprisingly, the numbers of calretinin-expressing neurons did not significantly change across developmental time points in surface fish (**Figure 3D**, left). In contrast, the numbers of cavefish calretinin neurons increased significantly across development and were greater in number than in surface fish (**Figure 3D**, left). The differences between surface and cavefish remained significant even when normalizing absolute numbers of calretinin neurons to OB volume. Therefore, the numbers of TH-expressing neurons increase broadly as a function of OB volume in both morphs, while calretinin neurons are selectively upregulated in cavefish (**Figure 3D**, right).

### Chemotopic organization in the OB is broadly conserved in both surface and cavefish

Different classes of odors evoke activity in different parts of the OB in *D. rerio* (Friedrich and Korsching, 1997, 1998; Li et al., 2005). We tested whether this chemotopic organization is conserved in *A. mexicanus* by performing *in vivo* Ca^2+^ imaging in transgenic lines of surface and cavefish expressing nuclear-localized GCaMP6s under the control of the elavl3 promoter (Tg(elavl3:GCaMP6s-H2B)) (Lloyd et al., 2022). The GCaMP6 expression patterns were qualitatively similar in the two morphs, with the somata of individual neurons visible in the baseline fluorescence, and glomerular regions of interest appearing dark presumably due to the nuclear-localization sequence (**Figure 4A**). We designed an 8-channel odor delivery system in which aquarium water with or without odor could be precisely delivered to the larval nares (**Figure 4B**). Each stimulus line was used for a single odor and were independently controllable via a solenoid valve. During imaging trials, water was constantly delivered to the nares from the water syringe. When the stimulus was triggered, the water valve closed and an odor (or control) valve opened.

**Figure 4:**
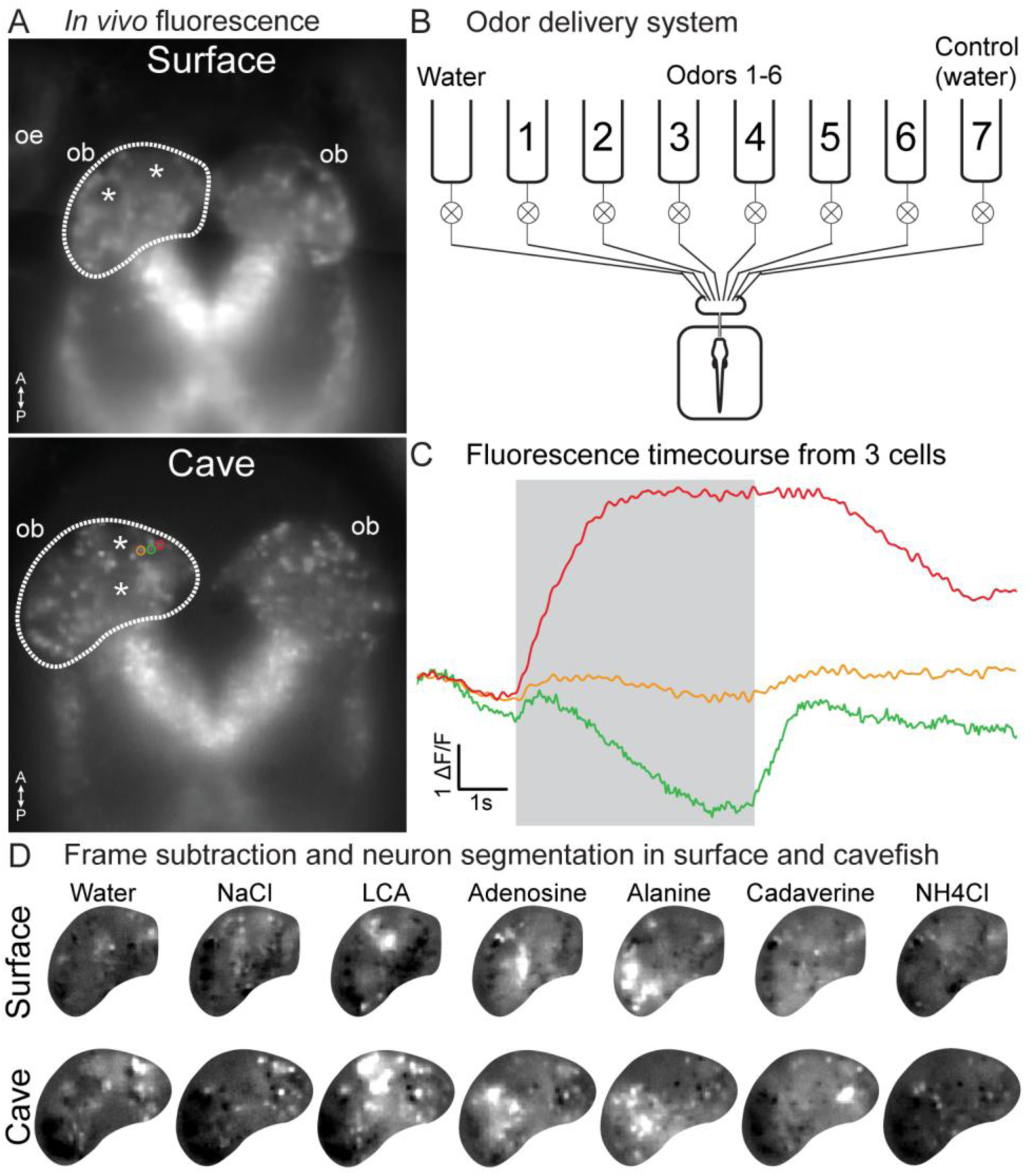
(A) *In vivo* mean fluorescence from surface (top) and cavefish (bottom) expressing soma-localized GCaMP6s. Asterisks indicate presumptive glomeruli in the OB. (B) The odor delivery system. (C) Odor responses from 3 cells from the cavefish example in panel (A). (D) Frame subtraction images from the preparations in panel (A) illustrating the spatial activity pattern evoked by 6 odor stimuli and water. Surface and cave examples are scaled to their own maximum. L, Lateral; M, Medial, A, Anterior; P, Posterior. OE, olfactory epithelium; OB, olfactory bulb.

**Figure 5:**
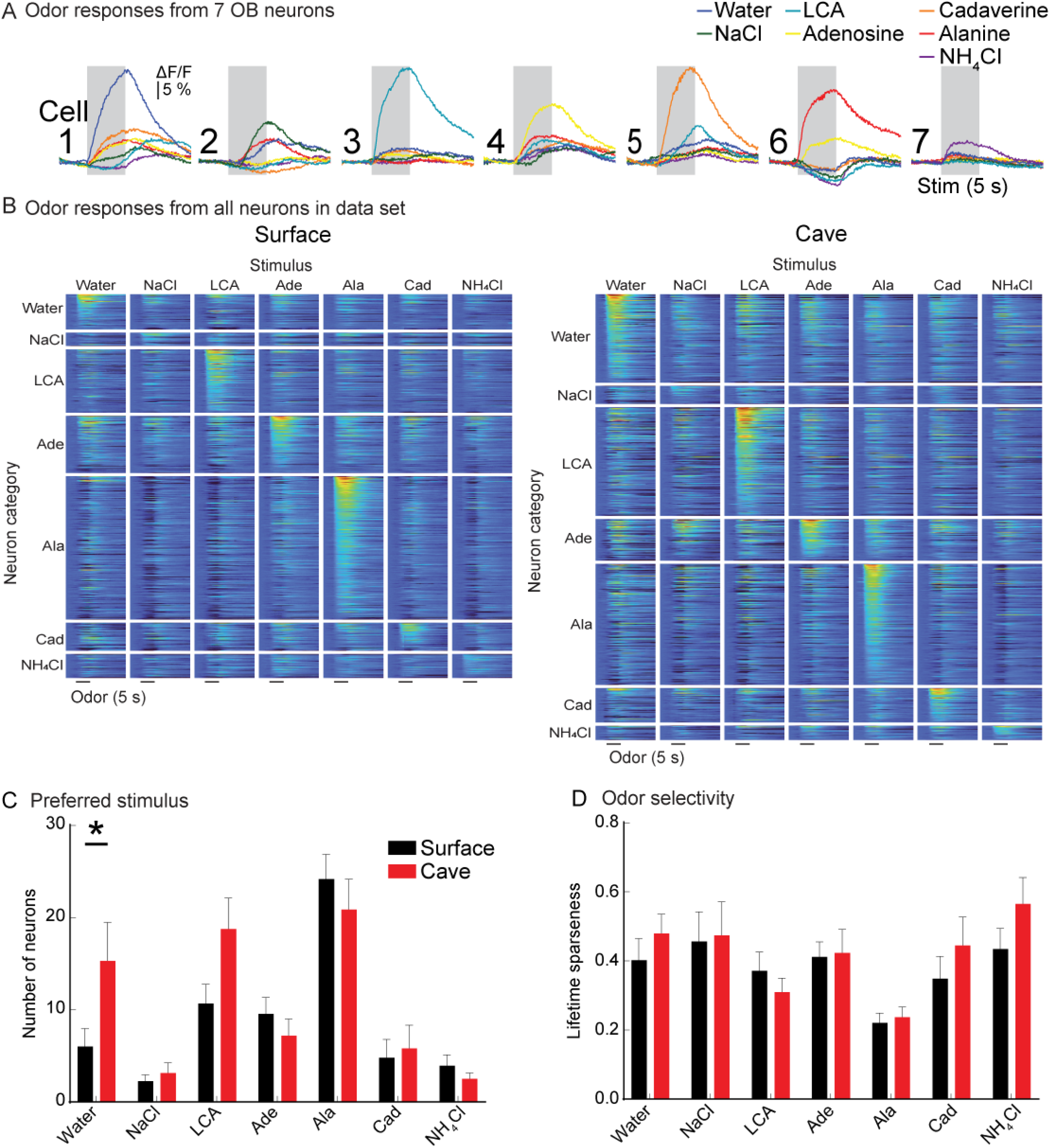
(A) Fluorescence time course from 7 different neurons in response to all tested odors. Each trace is the mean of 3 single trials. (B) Population heat map illustrating response of all neurons to each stimulus in the data set. Neurons are organized along the Y-axis first by their preferred odor, then by the strength of their response to that odor. (C) Number of neurons whose largest response was to each stimulus (“preferred” stimulus). (D) Mean lifetime sparseness of each neuron type.

We measured the response of OB neurons to a panel of odors known to be sensed by the olfactory system in other fish: the amino acid alanine, bile acid lithocholic acid (LCA), nucleic acid adenosine, sodium chloride (NaCl), cadaverine, ammonium chloride (NH_4_Cl), and a control water stimulus (Michel and Lubomudrov, 1995; Friedrich and Korsching, 1998; Herrera et al., 2021). Odors were delivered at concentrations between 10^-4^-10^-3^ M and evoked stimulus-locked increases and decreases in fluorescence that were detectable from individual neurons evident in the mean fluorescence, or in a frame subtraction analysis comparing the frames during and prior to the odor stimulus (**Figure 4C,D**).

Frame subtraction analysis of the spatial response to the different odors revealed the presence of odor-specific and spatially segregated peaks of activity across the OB (**Figure 4D**). Alanine strongly activated the lateral OB, LCA activated neurons in the medial OB, while the adenosine activated an intermediate region (**Figure 4D**). NaCl typically evoked a response that was medial to the LCA response, while cadaverine elicited a strong diffuse response, consistent with the activation of neurons outside of the imaging plane. NH_4_Cl produced a weak response, with a greater bias towards suppressed responses, and with no clear spatial organization. Notably, the “control” water stimulus activated a distinct set of neurons, with a core of activation in the most medial portion of the bulb. Qualitatively similar spatial patterns of activation were observed in all surface and cavefish morphs (**Figure 4D**).

### Cavefish exhibit significantly higher proportions of water-responsive OB neurons

All individual neurons that were detectable in the mean fluorescence or frame subtraction analysis were segmented for further analysis. Single trial measurements from 7 different neurons in response to the entire odor panel illustrate that most cells had a single stimulus which evoked the strongest response (**Figure 5A**). Each neuron was assigned to a “preferred” category according to the stimulus that evoked the largest response. The responses of all neurons within each category are illustrated in population heat maps sorted by the strength of their stimulus preference (**Figure 5B**, N = 16 surface fish, 61.4 +/- 5.51 SEM neurons per preparation; 16 cavefish; 73.6 +/- 9.3 SEM neurons per preparation). Surprisingly, all the odor-response categories had similar proportions of neurons in surface and cavefish, except for water-preferring neurons which were significantly more common in cavefish (surface: 6, cave: 15.3, *p*=0.035; 2-way repeated measures ANOVA: F_6,180_ = 2.57, *p=*0.02) (**Figure 5C**).

Each neuron exhibited variability in the degree of selectivity to the non-preferred odors; some responded primarily to a single odor, while others responded to multiple non-primary odors (**Figure 5A**, compare cell 3 to cell 4). We quantified the odor selectivity of individual neurons using the metric lifetime sparseness (LS), in which values near 0 indicate a selective neuron, while values near 1 indicate neurons that respond similarly to each stimulus. Alanine-preferring neurons were the most selective in both surface and cavefish (**Figure 5D**, surface: 0.2; cavefish: 0.22; REML: F_6, 135_ = 7.98, *p*<0.001; alanine comparison: p < 0.05 for all comparisons except LCA).

We further tested the result that OB neurons respond to water stimulation by measuring expression of the activity-dependent marker phosphorylated ERK (pERK). Surface and cavefish were paralyzed and mounted in agarose and were subjected to 5 minutes of repeated water stimulation from two lines alternating at a rate of 0.1 Hz. Control groups for both surface and cave morphs underwent the same process except the nares were not cleared of agarose prior to stimulation. The samples were then immediately euthanized, fixed, and underwent immunohistochemistry to label pERK-expressing cells (**Figure 6A**). Water stimulation in the experimental groups resulted in a significant increase in the number of pERK-expressing neurons in cavefish, but not surface fish (**Figure 6A**) (two-way ANOVA: F_1, 24_ = 7.06, p=0.014, Sidak’s post-hoc test, cavefish ctrl vs. stim: p=0.071). In both surface and cavefish, pERK labeled neurons were present throughout the entire OB, and did not co-localize with calretinin (not shown, n = 12 surface fish; n = 16 cavefish). Therefore, the OB of both surface and cavefish contain neurons that respond to water, although the cavefish OB contains significantly more water-responsive neurons.

**Figure 6:**
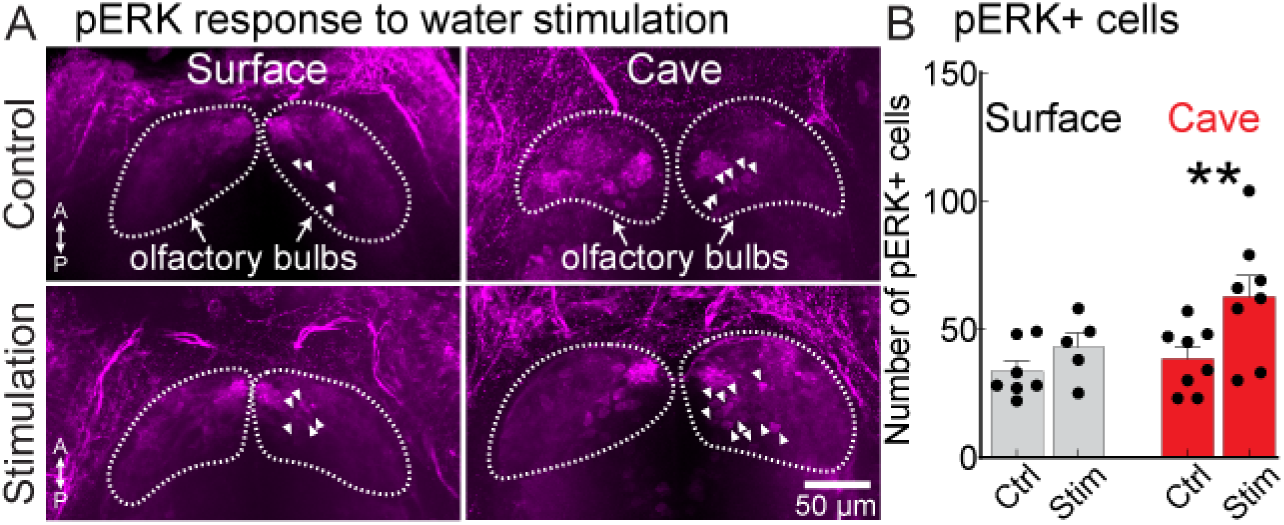
(A) Phosphorylated ERK (pERK) expression in the dorsal OB of surface and cavefish in response to water stimulation in a control group (nares blocked), and in an experimental group (nares exposed). The confocal images are maximum-projection from sections through the dorsal OB. Arrows denote pERK-positive neurons. (B) Number of pERK+ cells in the OB of surface and cavefish in control and stimulus groups. **, p < 0.01.

### The olfactory epithelium contains water-sensitive neurons that express Piezo2

We tested whether sensitivity to water might originate from the OE by imaging GCaMP6s-expressing neurons located in the OE in response to repeated presentations of water (**Figure 7**). Each stimulus evoked Ca^2+^ transients that originated from a subset of the neurons in the OE in both surface and cavefish (**Figure 7A-C**). We reasoned that the larger number of water-responsive neurons present in cavefish might reflect the presence of different levels of mechanosensitive proteins. We tested this possibility by performing whole-mount immunohistochemistry on the OE for the mechanosensitive ion channel Piezo2. Both the surface and cavefish OE contained Piezo2-expressing cells, although cavefish had significantly more Piezo2-expressing cells (**Figure 7D-E**). Overlap between Piezo2-expressing cells and the neuron-specific Nissl stain NeuroTrace confirmed that Piezo2 expression is present in neurons in the olfactory epithelium (**Figure 7F**). Therefore, cavefish exhibit enhanced integration of mechanosensory stimuli in the olfactory system.

**Figure 7.**
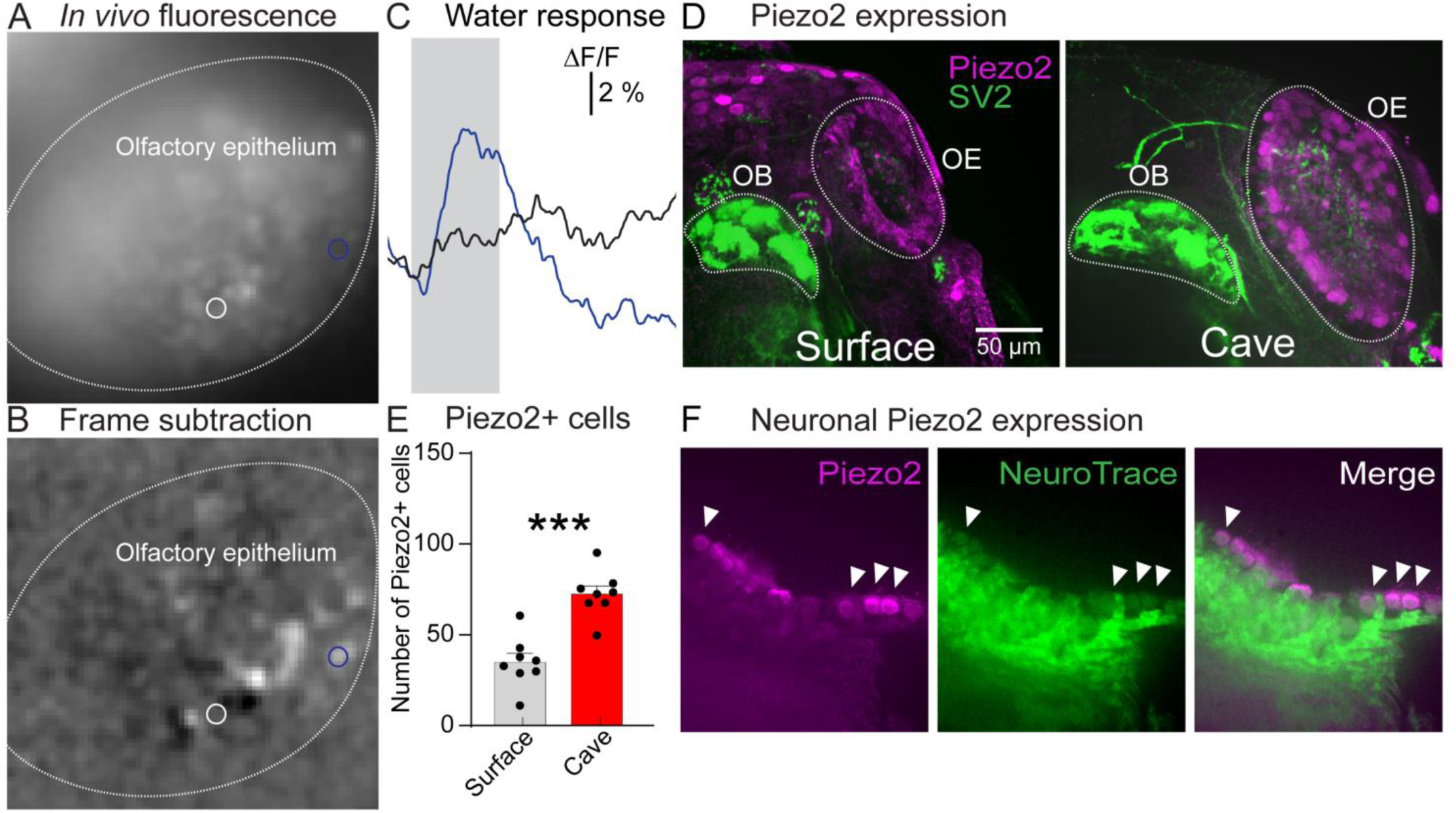
(A) Mean fluorescence of a elavl3:H2B-GCaMP6s transgenic cavefish. (B) Frame subtraction of (A) during stimulation with water. (C) Fluorescence time course from two cells in response to water stimulus at a flow rate of ∼300 µl min^-1^. The traces are from the regions of interest indicated in panels A-B. (D) Maximum intensity projection of the surface and cavefish OB immunoreacted for Piezo2 and SV2. (E) Number of Piezo2+ cells in the OE of surface and cavefish at 9 dpf. (F) OE of a cavefish co-labeled with Piezo2 immunofluorescence and NeuroTrace (arrows indicate cells with overlap). ***, p < 0.001.. OE, olfactory epithelium; OB, olfactory bulb.

**Table 1:** Statistical comparisons.

| Figure 1D - OB size | Surface | Cave | Stats |
| --- | --- | --- | --- |
| 7dpf | $1.91 \cdot 10^6 \mu\text{m}^3$ | $1.37 \cdot 10^6 \mu\text{m}^3$ | $t(24)=1.574, p=0.1286$ |
| **14dpf (\*\*)** | **$1.79 \cdot 10^6 \mu\text{m}^3$** | **$2.40 \cdot 10^6 \mu\text{m}^3$** | **$t(34)=3.383, p=0.0036$** |
| **21dpf (\*\*)** | **$1.81 \cdot 10^6 \mu\text{m}^3$** | **$3.43 \cdot 10^6 \mu\text{m}^3$** | **$t(9)=5.756, p=0.0008$** |
| **28dpf (\*\*\*)** | **$5.17 \cdot 10^6 \mu\text{m}^3$** | **$1.03 \cdot 10^7 \mu\text{m}^3$** | **$t(21)=4.611, p=0.0006$** |
| Figure 1E - ON diameter | Surface | Cave | Stats |
| 7dpf | 18.36 $\mu\text{m}$ | 16.99 $\mu\text{m}$ | $t(21)=0.717, p=0.4814$ |
| 14dpf | 17.79 $\mu\text{m}$ | 21.85 $\mu\text{m}$ | $t(11)=2.799, p=0.0510$ |
| 21dpf | 28.13 $\mu\text{m}$ | 31.23 $\mu\text{m}$ | $t(12)=2.298, p=0.0791$ |
| **28dpf (\*\*\*)** | **32.70 $\mu\text{m}$** | **58.49 $\mu\text{m}$** | **$t(9)=13.54, p<0.0001$** |
| Figure 2C (dlG) | Surface | Cave | Stats |
| 7dpf | $7.71 \cdot 10^3 \mu\text{m}^3$ | $7.96 \cdot 10^3 \mu\text{m}^3$ | $t(10)=0.2256, p=0.8261$ |
| 14dpf | $4.21 \cdot 10^4 \mu\text{m}^3$ | $5.13 \cdot 10^4 \mu\text{m}^3$ | $t(6)=1.351, p=0.5353$ |
| 21dpf | $5.37 \cdot 10^4 \mu\text{m}^3$ | $1.43 \cdot 10^5 \mu\text{m}^3$ | $t(8)=1.302, p=0.5353$ |
| **28dpf (\*\*\*)** | **$1.26 \cdot 10^5 \mu\text{m}^3$** | **$2.43 \cdot 10^5 \mu\text{m}^3$** | **$t(17)=4.615, p=0.0010$** |
| Figure 2C (lG) | Surface | Cave | Stats |
| 7dpf | $1.31 \cdot 10^4 \mu\text{m}^3$ | $9.36 \cdot 10^3 \mu\text{m}^3$ | $t(10)=2.483, p=0.1235$ |
| 14dpf | $4.70 \cdot 10^4 \mu\text{m}^3$ | $6.75 \cdot 10^4 \mu\text{m}^3$ | $t(6)=2.280, p=0.1768$ |
| 21dpf | $5.58 \cdot 10^4 \mu\text{m}^3$ | $2.11 \cdot 10^5 \mu\text{m}^3$ | $t(8)=1.318, p=0.2240$ |
| 28dpf | $2.29 \cdot 10^5 \mu\text{m}^3$ | $3.70 \cdot 10^5 \mu\text{m}^3$ | $t(17)=1.931, p=0.1768$ |
| Figure 2C (mdG) | Surface | Cave | Stats |
| 7dpf | $9.01 \cdot 10^3 \mu\text{m}^3$ | $1.05 \cdot 10^4 \mu\text{m}^3$ | $t(10)=1.473, p=0.4313$ |
| 14dpf | $5.28 \cdot 10^4 \mu\text{m}^3$ | $4.91 \cdot 10^4 \mu\text{m}^3$ | $t(6)=1.212, p=0.4689$ |
| 21dpf | $5.07 \cdot 10^4 \mu\text{m}^3$ | $1.04 \cdot 10^5 \mu\text{m}^3$ | $t(8)=1.073, p=0.4689$ |
| **28dpf (\*)** | **$1.05 \cdot 10^5 \mu\text{m}^3$** | **$1.68 \cdot 10^5 \mu\text{m}^3$** | **$t(17)=3.351, p=0.0151$** |
| Figure 2C (vmG) | Surface | Cave | Stats |
| 7dpf | $1.11 \cdot 10^4 \mu\text{m}^3$ | $1.43 \cdot 10^4 \mu\text{m}^3$ | $t(10)=2.348, p=0.1174$ |
| 14dpf | $4.58 \cdot 10^4 \mu\text{m}^3$ | $3.68 \cdot 10^4 \mu\text{m}^3$ | $t(6)=0.829, p=0.5246$ |
| 21dpf | $4.21 \cdot 10^4 \mu\text{m}^3$ | $1.61 \cdot 10^5 \mu\text{m}^3$ | $t(8)=1.083, p=0.5246$ |
| **28dpf (\*\*\*)** | **$1.46 \cdot 10^5 \mu\text{m}^3$** | **$3.40 \cdot 10^5 \mu\text{m}^3$** | **$t(17)=3.558, p=0.0096$** |
| Figure 2C (dG) | Surface | Cave | Stats |
| 7dpf | $3.58 \cdot 10^3 \mu\text{m}^3$ | $4.92 \cdot 10^3 \mu\text{m}^3$ | $t(10)=2.258, p=0.1359$ |
| 14dpf | $2.57 \cdot 10^4 \mu\text{m}^3$ | $2.60 \cdot 10^4 \mu\text{m}^3$ | $t(6)=0.109, p=0.9169$ |
| 21dpf | $3.11 \cdot 10^4 \mu\text{m}^3$ | $1.17 \cdot 10^5 \mu\text{m}^3$ | $t(8)=1.169, p=0.4761$ |
| **28dpf (\*\*\*)** | **$7.24 \cdot 10^4 \mu\text{m}^3$** | **$1.36 \cdot 10^5 \mu\text{m}^3$** | **$t(17)=4.673, p=0.0009$** |
| Figure 2C (maG) | Surface | Cave | Stats |
| 7dpf | $6.41 \cdot 10^3 \mu\text{m}^3$ | $5.66 \cdot 10^3 \mu\text{m}^3$ | $t(10)=1.318, p=0.6292$ |
| 14dpf | $3.26 \cdot 10^4 \mu\text{m}^3$ | $2.88 \cdot 10^4 \mu\text{m}^3$ | $t(6)=0.719, p=0.6292$ |
| 21dpf | $4.90 \cdot 10^4 \mu\text{m}^3$ | $9.87 \cdot 10^4 \mu\text{m}^3$ | $t(8)=0.997, p=0.6292$ |
| **28dpf (\*)** | **$1.16 \cdot 10^5 \mu\text{m}^3$** | **$3.72 \cdot 10^5 \mu\text{m}^3$** | **$t(17)=3.052, p=0.0286$** |
| Figure 2C (vpG) | Surface | Cave | Stats |
| 7dpf | $9.08 \cdot 10^2 \mu\text{m}^3$ | $1.62 \cdot 10^3 \mu\text{m}^3$ | $t(10)=1.319, p=0.4644$ |
| 14dpf | $3.90 \cdot 10^3 \mu\text{m}^3$ | $4.27 \cdot 10^3 \mu\text{m}^3$ | $t(4)=0.126, p=0.9061$ |
| 21dpf | $3.71 \cdot 10^3 \mu\text{m}^3$ | $7.73 \cdot 10^3 \mu\text{m}^3$ | $t(4)=1.586, p=0.4644$ |
| **28dpf (\*\*)** | **$2.47 \cdot 10^4 \mu\text{m}^3$** | **$5.36 \cdot 10^4 \mu\text{m}^3$** | **$t(17)=4.036, p=0.0034$** |
| Figure 2D (dlG norm.) | Surface | Cave | Stats |
| 7dpf | 0.2 | 0.1861 | $t(10)=0.605, p=0.7496$ |
| 14dpf | 0.2073 | 0.2417 | $t(6)=1.091, p=0.8021$ |
| 21dpf | 0.227 | 0.2326 | $t(8)=0.110, p=0.9928$ |
| 28dpf | 0.1807 | 0.1805 | $t(17)=0.008, p=0.9936$ |
| Figure 2D (lG norm.) | Surface | Cave | Stats |
| **7dpf (\*)** | **0.3353** | **0.2208** | **$t(10)=4.319, p=0.0106$** |
| 14dpf | 0.2632 | 0.2318 | $t(6)=3.543, p=0.0822$ |
| 21dpf | 0.2273 | 0.3051 | $t(8)=2.510, p=0.2285$ |
| 28dpf | 0.3188 | 0.2595 | $t(17)=1.2128, p=0.7499$ |
| Figure 2D (mdG norm.) | Surface | Cave | Stats |
| 7dpf | 0.2338 | 0.248 | $t(10)=1.129, p=0.5857$ |
| 14dpf | 0.2632 | 0.2318 | $t(6)=1.196, p=0.8022$ |
| 21dpf | 0.2419 | 0.1703 | $t(8)=1.687, p=0.5282$ |
| 28dpf | 0.1478 | 0.1256 | $t(17)=0.008, p=0.9936$ |
| Figure 2D (vmG norm.) | Surface | Cave | Stats |
| 7dpf | 0.2910 | 0.3407 | $t(10)=1.691, p=0.4775$ |
| 14dpf | 0.2172 | 0.1722 | $t(6)=1.415, p=0.7511$ |
| 21dpf | 0.1766 | 0.1984 | $t(8)=0.544, p=0.9367$ |
| 28dpf | 0.2042 | 0.2289 | $t(17)=0.9418, p=0.8321$ |
| Figure 2D (dG norm.) | Surface | Cave | Stats |
| 7dpf | 0.093 | 0.1157 | $t(10)=2.244, p=0.2588$ |
| 14dpf | 0.1248 | 0.1235 | $t(6)=0.118, p=0.9098$ |
| 21dpf | 0.1358 | 0.1585 | $t(8)=1.137, p=0.7435$ |
| 28dpf | 0.1043 | 0.09993 | $t(17)=0.344, p=0.9814$ |
| Figure 2D (maG norm.) | Surface | Cave | Stats |
| 7dpf | 0.124 | 0.105 | $t(10)=1.494, p=0.5162$ |
| 14dpf | 0.1584 | 0.1355 | $t(6)=1.125, p=0.8022$ |
| 21dpf | 0.2095 | 0.1579 | $t(8)=1.753, p=0.5282$ |
| 28dpf | 0.1643 | 0.2470 | $t(17)=1.940, p=0.3943$ |
| Figure 2D (vpG norm.) | Surface | Cave | Stats |
| 7dpf | 0.0174 | 0.0294 | $t(10)=1.209, p=0.5857$ |
| 14dpf | 0.0095 | 0.0229 | $t(6)=0.935, p=0.8021$ |
| 21dpf | 0.008901 | 0.009722 | $t(8)=0.11, p=0.9928$ |
| 28dpf | 0.03699 | 0.03907 | $t(17)=0.204, p=0.9814$ |
| Figure 3C (TH cells) | Surface | Cave | Stats |
| 7dpf | 82.71 | 111.1 | $t(12)=2.045, p=0.0634$ |
| **14dpf (\*\*\*)** | **46.44** | **100.8** | **$t(19)=4.789, p=0.0005$** |
| **21dpf (\*\*)** | **103.8** | **193.8** | **$t(9)=4.797, p=0.0029$** |
| **28dpf (\*\*)** | **284.8** | **484.0** | **$t(7)=4.301, p=0.0071$** |
| Figure 3C (TH cells norm.) | Surface | Cave | Stats |
| 7dpf | $6.00 \times 10^{-5}$ | $8.50 \times 10^{-5}$ | $t(12)=2.031, p=0.1828$ |
| **14dpf (\*\*)** | **$2.11 \times 10^{-5}$** | **$3.81 \times 10^{-5}$** | **$t(19)=3.761, p=0.0053$** |
| 21dpf | $6.13 \times 10^{-5}$ | $5.15 \times 10^{-5}$ | $t(9)=1.261, p=0.4208$ |
| 28dpf | $5.04 \times 10^{-5}$ | $4.45 \times 10^{-5}$ | $t(7)=0.885, p=0.4208$ |
| Figure 3D (CR cells) | Surface | Cave | Stats |
| 7dpf | 12.80 | 18.29 | $t(10)=1.195, p=0.2595$ |
| **14dpf (\*\*\*)** | **7.13** | **20.29** | **$t(13)=8.652, p<0.0001$** |
| **28dpf (\*\*)** | **18.75** | **81.83** | **$t(12)=4.418, p=0.0017$** |
| Figure 3D (CR cells norm.) | Surface | Cave | Stats |
| 7dpf | $7.00 \times 10^{-6}$ | $1.53 \times 10^{-5}$ | $t(10)=1.948, p=0.0800$ |
| **14dpf (\*\*)** | **$5.61 \times 10^{-6}$** | **$1.08 \times 10^{-5}$** | **$t(13)=3.844, p<0.0061$** |
| **28dpf (\*)** | **$4.4 \times 10^{-6}$** | **$8.67 \times 10^{-6}$** | **$t(12)=3.203, p=0.0151$** |
| Figure 5C (stim. pref.) | Surface | Cave | Stats |
| **2-way RM ANOVA (\*)** |  |  | **$F(6,180)=2.575, p=0.0204$** |
| **Water (\*)** | **6.00** | **15.31** | **$t(210)=2.836, p=0.0346$** |
| NaCl | 2.25 | 3.13 | $t(210)=0.266, p>0.9999$ |
| LCA | 10.69 | 18.75 | $t(210)=2.455, p=0.0988$ |
| Adenosine | 9.56 | 7.19 | $t(210)=0.723, p=0.9883$ |
| Cadaverine | 4.81 | 5.81 | $t(210)=0.305, p>0.9999$ |
| Alanine | 24.19 | 20.88 | $t(210)=1.001, p=0.9287$ |
| NH<sub>4</sub>Cl | 3.94 | 2.50 | $t(210)=0.438, p=0.9995$ |
| Figure 5D (Odor selectivity) | Surface | Cave | Stats |
| **REML (\*\*\*)** |  |  | **$F(6,135)=7.978, p<0.0001$** |
| Water | 0.3868 | 0.4614 | $t(165)=0.9516, p=0.9470$ |
| NaCl | 0.4630 | 0.4916 | $t(165)=0.3249, p>0.9999$ |
| LCA | 0.3633 | 0.3098 | $t(165)=0.6997, p=0.9904$ |
| Adenosine | 0.4037 | 0.4176 | $t(165)=0.1739, p>0.9999$ |
| Cadaverine | 0.3206 | 0.4553 | $t(165)=1.537, p=0.6109$ |
| Alanine | 0.2205 | 0.2366 | $t(165)=0.2125, p>0.9999$ |
| NH<sub>4</sub>Cl | 0.4311 | 0.5607 | $t(165)=1.481, p=0.6538$ |
| Figure 6B (pERK+ cells) | Ctrl. | Stim. | Stats |
| **2-way RM ANOVA (\*)** |  |  | **$F(1,24)=7.062, p=0.0138$** |
| Surface | 33.57 | 43 | $t(24)=0.982, p=0.559$ |
| **Cave (\*\*)** | **38.5** | **62.63** | **$t(24)=2.942, p=0.0142$** |
| Figure 7B | Surface | Cave | Stats |
| **Piezo2+ cells** | **35.13** | **73** | **$t(14)=5.618, p<0.0001$** |

## Discussion

Here we provide the first data describing the structure and function of the first stage of olfactory sensory processing in *A. mexicanus* surface and blind cave morphs. Both surface and cave morphs share many general organizational and functional features with other teleosts, which includes a general glomerular organization in the OB, the presence of multiple interneuron cell types, and functional responses to a range of ethologically relevant chemical stimuli within

However, we observed several striking differences across the surface and cavefish morphs. This included a prominently larger OB, which contained larger glomerular regions of interest and increased numbers of interneurons within the OB (**Figures 1-3**). The most striking difference was the presence of more neurons within the cavefish OB exhibiting sensitivity to water, which was the only stimulus within our panel exhibiting a significant difference (**Figure 4-6**). We also confirmed that the OE of both surface and cavefish contain water-responsive neurons and neurons that express the mechanosensitive ion channel Piezo2 (**Figure 7**). Notably, the cavefish OE contained significantly greater numbers of Piezo2-expressing cells (**Figure 7**). Therefore, cavefish exhibit enhanced multisensory integration in the first stages of olfactory sensory processing, demonstrating the presence of a constructive sensory adaptation in cavefish. We propose that an increase in mechanical sensitivity may facilitate the ability of cavefish to detect and track behaviorally relevant olfactory cues in the cave environment.

### Conserved structure and function of the olfactory system

We observed a high degree of structural similarity between the OB of developing *A. mexicanus* and published data on zebrafish, and all 7 glomerular regions which are present in developing zebrafish have equivalent structures in both surface and cave *A. mexicanus* (Braubach et al., 2012; Braubach et al., 2013). This was also true of our functional imaging data, which showed responses to the amino acid alanine in the lateral region, the nucleotide adenosine in the intermediate region, and the bile acid LCA in the medial region, consistent with zebrafish imaging studies of olfactory response (Friedrich and Korsching, 1997; Li et al., 2005). Taken together, these results show that *A. mexicanus* exhibit broadly conserved chemotopy similar to what has been reported in *D. rerio*, and that these responses are relatively conserved across the two morphs.

### Broad changes in size of the olfactory system

It is thought that nutrient scarcity has been a primary driver of evolution in this species, and so studies of olfaction in *A. mexicanus* have largely been focused on increased sensitivity and attraction to nutrient stimuli (Hüppop, 1987; Bibliowicz et al., 2013; Blin et al., 2018; Blin et al., 2020; Blin et al., 2024). In principle, uniform increases in glomerular volume would reflect a broad increase in the number of overall OSNs, while selective or disproportionate increases in some glomeruli might reflect changes in specific subsets of OSNs. For example, *D. sechellia*, which subsists almost entirely on the noni fruit *Morinda citrifolia*, exhibits both an increase in the number of noni-sensitive Or22a-receptor neurons, and a concomitant increase in volume of the target glomeruli, relative to closely related species *D. melanogaster* (Dekker et al., 2006; Auer et al., 2020). We observed that all glomerular regions of interest increased proportionally with the size of the OB, suggesting a relatively uniform increase in input from different OSNs. Additionally, we did not observe gross changes in the population representation of the different amino acids in our odor panel (**Figure 5C**). These data are consistent with a model in which evolution of cavefish olfaction occurred in a broad manner and is not specifically geared towards enhanced detection of nutrient stimuli. However, we could not evaluate differences in detection threshold in this study as we only used a single concentration of each odor. Additionally, differential changes in sensitivity could also reflect differences in expression levels of olfactory receptors within OSNs, which would not result in changes in glomerular volume. Future studies are needed to determine whether surface and cave morphs exhibit different olfactory thresholds to the panel of odors used in our study.

### Increases in interneuron populations

We found increases in cell number of both dopaminergic and calretinin-positive interneurons in the cavefish OB. DA neurons were present in both the glomerular layer and the mitral cell layer, as in other teleost models (Byrd and Brunjes, 1995; Kawai et al., 2012). Dopamine interneurons have complex roles in both odor detection and discrimination, stemming from the action of DA and GABA on different cellular targets (Bundschuh et al., 2012; Lyons-Warren et al., 2023). Several studies have suggested that changes in dopamine synthesis, expression, or signaling, may contribute to cavefish morphological or behavioral adaptations (Elipot et al., 2014; Gallman et al., 2020; Jaggard et al., 2020). A leading hypothesis for the repeated evolution of albinism in cavefish is that the increased availability of L-tyrosine that would otherwise be used for melanin synthesis enables increased catecholamine synthesis, which may be broadly adaptive for cavefish (Bilandžija et al., 2013; Bilandžija et al., 2018; O’Gorman et al., 2021). Future studies investigating the relationship between OB interneuron populations and pigmentation would shed light on whether DA expression in the OB is linked with pigmentation levels.(Klaassen et al., 2018).

Calretinin neurons have also been reported in the teleost and mammalian olfactory bulb, but their distribution appears to vary across species (Porteros et al., 1997; Díaz-Regueira and Anadón, 2000; Castro et al., 2006; Parrish-Aungst et al., 2007; Castro et al., 2008). In mice and zebrafish, calretinin cells are found in the glomerular layer (Castro et al., 2006; Parrish-Aungst et al., 2007), however we found Calretinin-positive cell bodies almost exclusively in the dorsomedial region of the glomerular layer. This may be due to differences in the developmental time point studied, as we focused on the first month of development, and calretinin cells are known to be continuously generated throughout life (Batista-Brito et al., 2008). It is notable that the location of Calretinin-positive cell bodies appears to coincide with the location of mechanical responses in the bulb, which were also increased in cavefish. Calretinin neurons in the bulb are thought to function in a primarily intra-glomerular fashion, to filter out random excitation from the olfactory nerve and improve signal-to-noise ratio (Iseppe et al., 2016; Capsoni et al., 2021). This could be related to their expression changes in the mediodorsal region of the OB, where increased input from mechanosensitive neurons could necessitate increased inhibitory control. Further studies are needed to determine what, if any, role calretinin neurons play in mechanical responses in the bulb.

### Functional relevance of Piezo2-expression in the OE

The result that the cavefish OE contains significantly more Piezo2-expressing cells is consistent with the possibility that OB water sensitivity originates peripherally rather than through centrifugal inputs from non-olfactory sensory systems. Future studies are needed confirming the cell types that express Piezo2 in the OE and whether it is expressed in specific OSN subtypes. Indeed, this possibility is supported by our observation that water evoked a core of activation in the dorsomedial region of the OB (**Figure 4D**).

Although mechanosensitivity has not been described in the olfactory system of teleosts, the olfactory epithelium of *D. rerio* contains Piezo2 expression that is consistent with what we report here (**Figure 7**) (Aragona et al., 2024). Mechanical sensitivity is a well-established phenomenon in the olfactory system of both vertebrate and invertebrate brains. For example, a subset of olfactory receptors exhibit intrinsic mechanosensitivity (Grosmaitre et al., 2007; Connelly et al., 2015), and similar results have been described in mice and insects (Iwata et al., 2017; Mahajan et al., 2025). These processes may potentiate responses to weak odor activation or aid in odor tracking (Tiraboschi et al., 2021; Tuckman et al., 2021). Future studies investigating Piezo2 function in the OB could shed light on the role of these proteins in mechanical sensitivity of the olfactory system.

### Methodological considerations

The transgenic fish used in our study express GCaMP6s under control of the elavl3 promoter, which reportedly drives soma-localized expression in many cell types (Park et al., 2000; Freeman et al., 2014). Therefore, the functional imaging results reflect measurements from a diverse population of cell types that includes interneurons and projection neurons. Different OSNs map to OB glomeruli, which synapse onto neurons connected to that glomerulus. Therefore, the spatial localization of the functional signals likely reflect the postsynaptic activity of the OSNs innervating nearby glomeruli.

Our conclusions that cavefish exhibit enhanced mechanosensory responses are based on the observation that their OB contains greater numbers of neurons that respond to our water control stimulus. Our olfactometer is designed to provide a constant stream of aquarium water, which is then switched to one of seven lines containing the different stimuli (i.e., the same water with or without odor). The switching takes place via the opening and closing of two solenoid valves, a process which takes place in ∼3 msec (NResearch), and which will necessarily introduce a brief perturbation in the water flow. In principle, this kind of change could be detected by cells expressing the mechanosensitive ion channel Piezo2 (Woo et al., 2014).

### Conclusions

*A. mexicanus* provide a unique opportunity to study the genetic and neural basis of sensory evolution, due to their starkly different sensory phenotypes, recent divergence times, and amenability to laboratory study. Although here we studied only surface and cavefish descended from the Pachón cave population, there are over 30 caves in central Mexico harboring cave populations of *A. mexicanus*, with at least two completely independent lineages of cave fish descendent from surface forms of *A. mexicanus* (Herman et al., 2018; Moran et al., 2023). Indeed recent work examining the olfactory system in Molino cavefish indicates the presence of parallel adaptations (Harkinish-Murray et al., 2025). Future studies using this unique model organism will facilitate our understanding of how evolution drives adaptation and will provide insight into the underlying principles and organization of the brain.

## Acknowledgements

This work was supported by funding from DC000044, DC020519, and FSU’s Vice President for Research Scholarship. We are grateful to the Keene lab for generously sharing the *A. mexicanus* strains used in this study, and to members of the Storace laboratory for helpful discussion.

